# Ethyl-iophenoxic acid as a serum biomarker for marsupial species in oral bait trials

**DOI:** 10.64898/2026.03.13.711545

**Authors:** Sally A. Nofs, Ruth Pye, David Nichols, Shylo R. Johnson, Amy T. Gilbert, Billie Lazenby, Andrew S. Flies

## Abstract

Ethyl-iophenoxic acid (Et-IPA) is widely recognized as a useful biomarker to confirm oral bait consumption in eutherian species. In historical studies on marsupials, Et-IPA was rapidly eliminated from brushtail possums (*Trichosurus vulpecula*) and swamp wallabies (*Wallabia bicolor*) suggesting limited use for marsupial species. However, a 1 mg oral dose of Et-IPA was detectable in the marsupial Tasmanian devils (*Sarcophilus harrisii*) for ≥ 56 days suggesting the biomarker can be used in a devil bait vaccine program. To assess Et-IPA marking in off-target marsupials that may consume baits, we administered 1 mg oral doses of Et-IPA to brushtail possums, forester kangaroos (*Macropus giganteus tasmaniensis*), spotted-tailed quolls (*Dasyurus maculatus*) and eastern quolls (*Dasyurus viverrinus*). Liquid chromatography with tandem mass spectrometry was used to detect and quantify serum Et-IPA. Et-IPA was detected in the serum on day 2 but was not detected by day 14 in any of the species tested, including the two quoll species which are in the same carnivorous Dasyuridae family as the devils. The rapid elimination of Et-IPA in the marsupials included in this study suggests it is not useful as a biomarker for these species. Furthermore, rapid elimination in the kangaroos and possums suggests that Et-IPA is unlikely to accumulate in the food chain following distribution of Et-IPA-marked oral bait vaccines for Tasmanian devils.

**Short summary for non-experts:** A recent study in Tasmanian devils (*Sarcophilus harrisii*) challenged the concept that ethyl iophenoxic acid (Et-IPA) is not a useful serum biomarker for marsupials. Using the same sensitive liquid chromatography-tandem mass spectrometry method we detected serum Et-IPA in four marsupial species on day two post-ingestion but by day 14, serum Et-IPA was undetectable. These findings indicate that Et-IPA is an unsuitable biomarker for these species and suggest that Et-IPA from devil bait vaccines is unlikely to bioaccumulate in the Tasmanian environment.

## Introduction

Oral baits are effective tools in wildlife management for the delivery of various xenobiotics including vaccines. Biomarkers in baits enable the evaluation of bait uptake by target and off-target species to determine the success of a bait distribution strategy. Various biomarkers have been used in oral baiting systems including rhodamine B, tetracycline, and derivatives of iophenoxic acid as reviewed in Fry and Dunbar, 2007. While some authors report problems with palatability of rhodamine B, this is an effective bait marker for eutherian and some marsupial species including common brushtail possums (*Trichosurus vulpecula*), tamar wallabies (*Notamacropus eugenii*) and northern quolls (*Dasyurus hallucatus)* (Morgan, 1981, Williams, 1997, Heiniger et al., 2018).

Ethyl-iophenoxic acid (Et-IPA) (a-Ethyl-3-hydroxy-2,4,6-triiodohydrocinnamic acid) has been used effectively in eutherian mammals as a bait marker, as reviewed in Ballesteros et al., 2013, yet studies in the 1990’s suggested it was of little use in marsupial species. A comparative study between domestic cats (*Felis catus*) and common brushtail possums found the plasma elimination half-life of Et-IPA was 107 days for cats administered 1.5 mg Et-IPA/ kg bodyweight compared to one day in possums given either 1.5 mg/kg or 10 mg/kg (Eason et al., 1994). In these possums, serum Et-IPA returned to pre-treatment levels within five to ten days. A separate study on swamp wallabies (*Wallabia bicolor*) administered up to 30 mg Et-IPA /kg bodyweight found that serum Et-IPA levels returned to baseline by day six (Fisher and Marks, 1997). Both studies concluded that Et-IPA was unsuitable as a long-term biomarker in marsupial species, perhaps due to marsupials having a weaker plasma protein binding affinity for Et-IPA and/ or a different Et-IPA excretion pathway compared to eutherian mammals.

The method used for detection of serum Et-IPA in both these marsupial studies was the serum iodine assay, an indirect measurement of serum Et-IPA that measures protein-bound iodine. Liquid chromatography with mass spectrometry is a direct and more sensitive method for detection and quantification of the biomarker. Studies by Spurr, 2002 and Purdey et al., 2003 provide a comparison between the indirect and direct methods of serum Et-IPA detection in stoats (*Mustela erminea*). Spurr 2002 gave stoats an oral dose of between 3 and 17 mg/kg Et-IPA and reported to find elevated iodine for 14 days using plasma iodine detection. Purdey et al. 2003 administered 8 mg/kg Et-IPA to stoats and using high performance liquid chromatography, detected serum Et-IPA at day 27 when the trial ended. The authors discussed the benefits of liquid chromatography over the indirect serum iodine method for measuring serum Et-IPA which included less susceptibility to false negative results. Additionally, high-performance liquid chromatography resulted in Et-IPA detection in opossums (*Didelphis virginiana*) during a field application of Et-IPA-marked baits (Campbell et al., 2006).

An oral bait vaccine to protect Tasmanian devils against devil facial tumour disease (DFTD) is currently in development (Flies et al., 2020). Preliminary field evaluation of baiting strategies targeting Tasmanian devils revealed that the baits were often taken and consumed by off-target species (Dempsey et al., 2023). The omnivorous brushtail possum and the herbivorous macropod, Tasmanian pademelon (*Thylogale billardierii*), were noted to eat the meat-based baits in the placebo field trials. The carnivorous spotted-tailed quolls and eastern quolls have also been observed to readily eat the placebo baits (S. Nofs unpublished). The pilot baiting study reported that 93% of baits placed on the ground were consumed by off-target species (Dempsey et al., 2023). In the same study, use of a bait dispenser designed for raccoons in North America (Smyser et al., 2015) decreased off-target bait consumption to 45%.

Due to the challenges of delivering baits to wild devils, an effective bait biomarker is required to determine appropriate strategies for devil facial tumour bait vaccine distribution. Liquid chromatography with tandem mass spectrometry (LC-MS/MS) detected serum Et-IPA in devils for up to 56 days following consumption of a 1 mg dose (Pye et al., 2023). Doses of 50 mg per animal resulted in detection of serum Et-IPA beyond 200 days. The success of this study encouraged us to determine whether the sensitive LC-MS/MS method could detect Et-IPA in serum of other marsupial species for a longer period than had been previously reported.

Here we tested the serum biomarker capacity of a 1 mg oral dose of Et-IPA across four marsupial species: brushtail possums, forester kangaroos (*Macropus giganteus tasmaniensis*), spotted-tailed quolls (*Dasyurus maculatus*), and eastern quolls (*Dasyurus viverrinus*) representing omnivorous, herbivorous, and carnivorous marsupial species commonly found in Tasmania. We hypothesized that the sensitivity of the LC-MS/MS method would allow for the detection of serum Et-IPA in the brushtail possum and forester kangaroo for longer than reported in previous studies on marsupials that used the indirect serum iodine method of detection. We expected both species of quoll, in the same Dasyuridae family as the devil, to have similar results to the devils. Despite the increased sensitivity of the test, our results are in agreement with prior studies that Et-IPA is not a suitable biomarker for many marsupials.

## Methods

### Animals, and rationale for species selection

Two individuals from each of four species: common brushtail possums, forester kangaroos, spotted-tailed quolls and eastern quolls, were used in this study (Figure 1). The selection of individuals was dependent on resident animal availability at the cooperating institutions. Macropod inclusion in the study was carefully considered due to the risk of capture myopathy (McMahon et al., 2013, Vogelnest and Woods, 2008). Although forester kangaroos were not observed in the preliminary bait trials, likely because of their limited geographic range in Tasmania, this species was selected for our study as a surrogate to represent the Tasmanian macropods. The availability of forester kangaroos and their habituation to humans greatly facilitated Et-IPA administration, sedation and sample collection. The young adult forester kangaroos used were within the weight range of adult Bennett’s wallabies (*Notamacropus rufogriseus*), a common macropod species in Tasmania.

**Figure 1.**
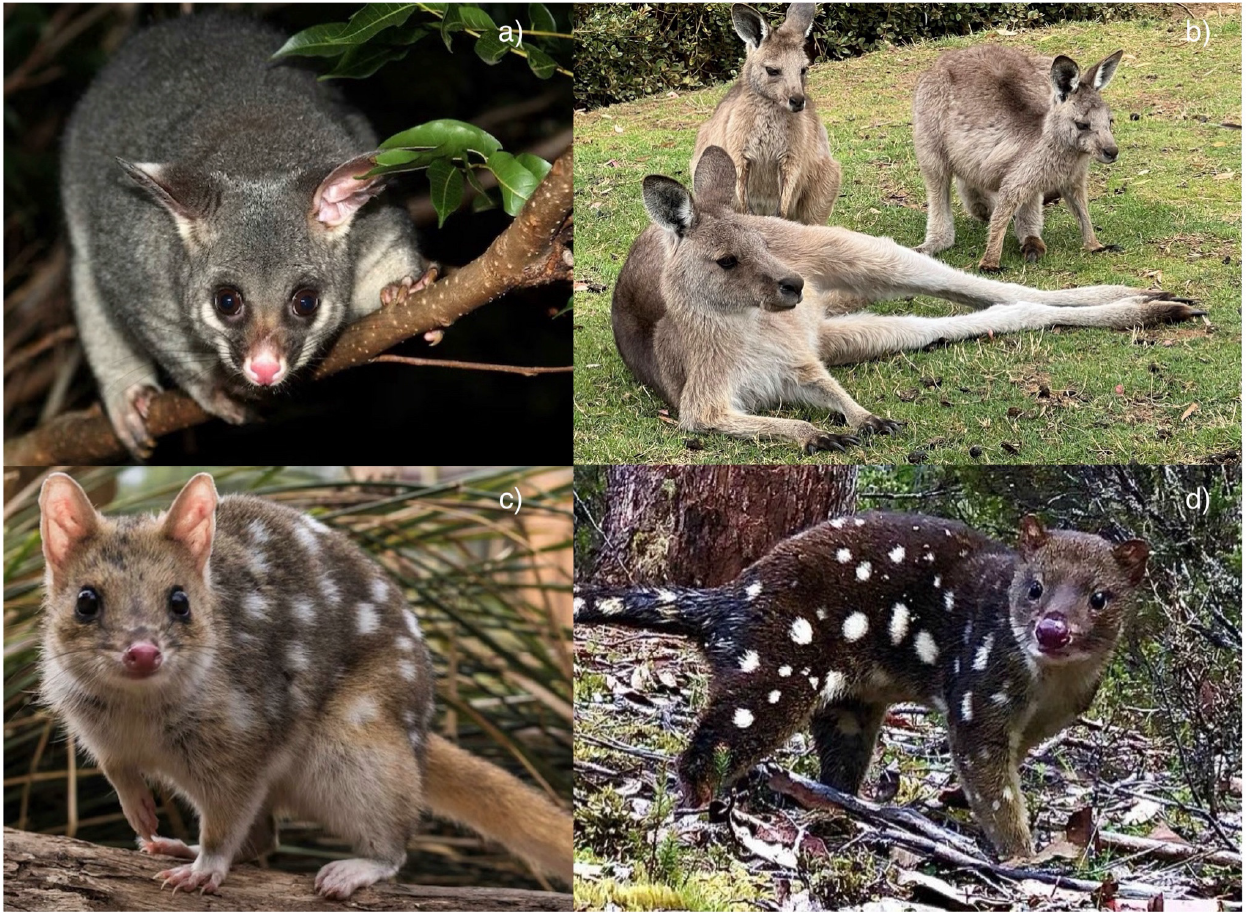
Marsupial species included in this study: a) common brushtail possum, b) forester kangaroo (young adult in background), c) eastern quoll and d) spotted-tailed quoll. Photo credits/ supplied by: a) Brisbane City Council, b) Sally Nofs, c) Parks Australia, d) David Hamilton.

The eight individuals are listed in Table 1 along with their age, sex and body weight. All procedures were approved by the University of Tasmania Animal Ethics Committee under AEC#28506.

**Table 1.** Details of individual study animals. Species, study identification, sex, age (years), body weight (kg), dosage (mg/kg) Et-IPA, method of oral administration of Et-IPA, serum Et-IPA concentration in ng/ml on day 2 and day 14. All animals received 1 mg of Et-IPA.

| Species | Study ID & Last 4 digits of microchip | Sex | Age (yrs) | Weight (kg) | Et-IPA dose (mg/kg) | Vehicle for oral administration | Serum Et-IPA (ng/ml) Day 2 | Serum Et-IPA (ng/ml) Day 14 |
| --- | --- | --- | --- | --- | --- | --- | --- | --- |
| common brushtail possum | BTP1 2925 | M | 4.0 | 4.2 | 0.24 | porridge* | 4.3 | 0** |
| common brushtail possum | BTP2 3841 | M | 1.5 | 3.06 | 0.33 | green grape | 20.0 | 0** |
| forester kangaroo | FK1 3680 | F | <1.5 | 9.1 | 0.11 | oral | 2.9 | 0** |
| forester kangaroo | FK2 3839 | F | <1.5 | 11.4 | 0.09 | oral | 4.0 | 0** |
| spotted-tailed quoll | STQ1 1164 | F | 2.6 | 2.7 | 0.37 | day-old chick | 399.0 | 0** |
| spotted- tailed quoll | STQ2 3833 | F | 2.6 | 3.4 | 0.29 | day-old chick | NA | 0** |
| eastern quoll | EQ1 3147 | F | 3.7 | 0.89 | 1.12 | meatball* | 317.7 | 0** |
| eastern quoll | EQ2 7697 | F | 3.7 | 0.9 | 1.1 | meatball* | 455.3 | 0** |

\*Ingredients in supplementary material
\*\* < Limit of Quantitation of 0.5 ng/ml
NA = not available

### Housing and husbandry

The possums were housed individually in enclosures measuring 2.4 m (W) x 2.3 m (H) x 7 m (L). Their diet was a mix of vegetables and fruit supplemented with native flowers, eggs, porridge and dog kibble. The kangaroos were managed as a social group of approximately 150 individuals in a 1.4 hectare paddock. Their diet included the paddock pasture, hay and macropod pellets. Spotted-tailed quolls were housed in pairs in an enclosure measuring 10 m (W) x 2 m (H) x 10.5 m (L). Eastern quolls were housed in pairs in a 5 m (W) x 2 m (H) x 10.5 m (L) enclosure. All quolls were fed commercially sourced carcasses and/ or off-cuts of a variety of species including wallaby (*Notamacropus rufogriseus*) and chicken (*Gallus gallus*). The eastern quolls were also fed supplemental ‘quoll balls’ based on minced chicken. All enclosures contained a variety of enrichment items including climbing logs and wooden boxes to mimic natural burrows. All animals had free access to fresh water.

### Et-IPA administration

Et-IPA (α-Ethyl-3-hydroxy-2,4,6-triiodohydrocinnamic acid, CAS 96-84-4, Sigma Aldrich) for oral administration was prepared as in the prior study for Tasmanian devils (Pye et al., 2023) and as adapted from an earlier reference (Ballesteros et al., 2013). Ten mg of Et-IPA powder was added to 1.0 mL edible corn oil, vortexed, then heated to 90°C to aid dissolution. Next, 100 µl of the 10 mg/mL solution was incorporated within each administration vehicle. At 1 mg Et-IPA per animal, the variable body weights resulted in dosage ranges for individuals from 0.09 mg/kg (forester kangaroo) to 1.1 mg/kg (eastern quoll), (Table 1).

A method for Et-IPA administration for each species (Table 1) was determined through consultation with animal care staff. Et-IPA was administered to one possum in a porridge mixture (recipe in supplemental material), and to the other possum in a fresh green grape. Kangaroos received the Et-IPA solution orally via syringe during a brief manual restraint, and spotted-tailed quolls were administered their doses in thawed day-old-chickens. All Et-IPA administration was directly observed to monitor consumption. Eastern quolls were shy, mainly nocturnal, and unlikely to eat during observation. They were therefore temporarily individually housed in pest-proof transport crates overnight and the Et-IPA dose was delivered in a meatball (recipe in supplemental material). For each quoll, the marked meatball was completely consumed by the morning. Due to the overnight bait consumption by the eastern quolls, blood collection was performed on day two following bait offering for all individuals.

### Animal capture, sedation or general anaesthesia, and blood sample collection

All animals used in this study were captured and then anaesthetised to enable blood sample collection. Following Et-IPA administration on day 0, serial blood sample collection was planned for days 2, 14, 28 and 56 depending on serum Et-IPA detection.

Possums were netted or manoeuvred into a crate or soft fabric sack. For both the spotted-tailed and eastern quolls, PVC pipe traps (approximately 315 mm diameter and 875 mm in length) developed for Tasmanian devils (Hawkins et al., 2006) were baited with meat and set up in enclosures, with nets used for capture if the traps were unsuccessful. Each animal was then transferred to a hessian or fabric sack. While in the sack, induction of anaesthesia occurred using 5% isoflurane and 1 to 2.5 L/ minute oxygen delivered through the fabric via a mask placed over the muzzle (Vogelnest and Woods, 2008). Once immobile, the animal was removed from the sack and the mask placed directly over the muzzle to maintain anaesthesia at 2% isoflurane and 1 to 2.5 L/minute oxygen.

The forester kangaroos were easily approached in their 1.4 ha enclosure and were sedated with an intramuscular injection of 50-60 ug/kg of medetomidine (1.0 mg/ml Medetate® medetomidine hydrochloride, Jurox, NSW, AU) (Dr. Evie Clark, personal communication). They were then observed until recumbent and unresponsive. This approach minimised stress and the risk of capture myopathy from restraint. On completion of procedures, atipamezole 5 mg/mL (atipamezole hydrochloride, Ilium, Troy Lab, NSW, AU), the reversal agent for medetomidine, was administered intramuscularly in the same volume as medetomidine and the kangaroos were observed closely until ambulatory, alert, and responsive. All animals recovered from anaesthesia and sampling procedures with no adverse effects.

All animals in the study were given physical examinations while anaesthetised, with microchips placed or existing microchips verified. A patch of fur was shaved from the kangaroos to aid visual identification. Blood (1 to 3 mL) was collected from the lateral tail vein (possums and kangaroos), or jugular or ventral tail vein (quolls) and immediately placed into clot activating tubes. Within four hours of collection, blood samples were centrifuged for five minutes at 1,700 x *g*, and serum pipetted into 1.5 mL cryotubes and stored at −70°C until analysis.

### Detection of serum Et-IPA and LC-MS/MS protocol

Baseline (pre-treatment) blood samples were not collected from the individual animals to minimize the number of anaesthetic events. Instead, archived serum samples opportunistically collected at the time of euthanasia from one brushtail possum, two forester kangaroos (pooled) and five eastern quolls (pooled) were used to construct the matrix-matched external calibration curves for sample quantitation of Et-IPA (Figure 2).

**Figure 2.**
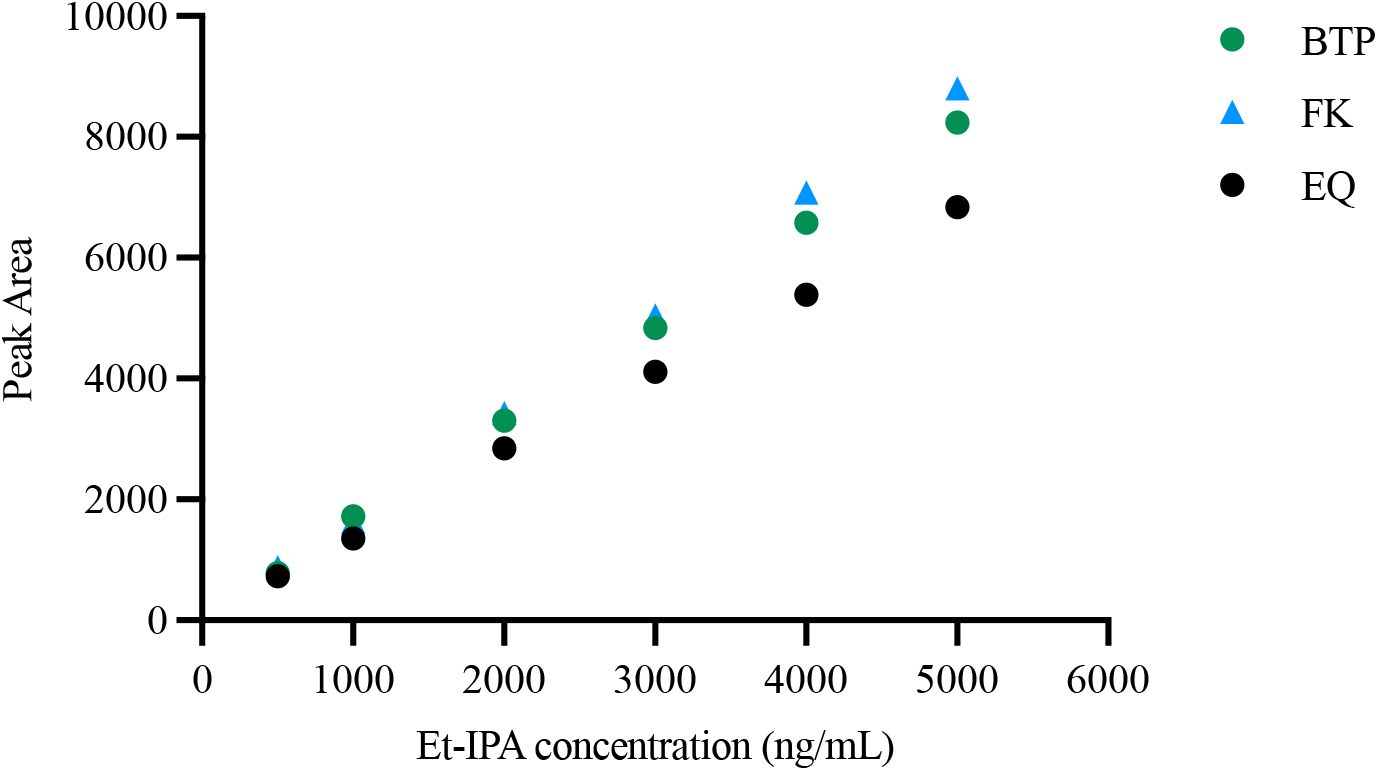
Serum calibration comparison for brushtail possum, forester kangaroo and eastern quoll demonstrating species-specific effects of serum as a sample matrix on calibration slopes increasing at higher Et-IPA concentration levels. BTP: brushtail possum, 1 serum sample; FK: Forester kangaroo, 2 x pooled serum samples; EQ, eastern quoll, 5 x pooled serum samples.

Once calibration was completed study serum samples were prepared as previously described (Pye et al., 2023). In brief, 200 µL of serum was mixed with 800 µL of Methanol: Acetone (1:1, v/v) then vortexed for 30 seconds. To encourage protein precipitation, the sample was then placed at −20°C for 30 minutes. After 12 minutes of centrifugation at 21 000 *g* and 4°C the supernatant was transferred to an ultra-performance liquid chromatography (UPLC) vial.

### Sample analysis

LC-MS/MS analytical methodology was the same as reported for Tasmanian devils (Pye et al., 2023). A Waters Acquity® H-class UPLC system (Waters Corporation, Milford, MA) analysed serum Et-IPA concentrations. A Waters Acquity VanGuard C18 pre-column (5 x 2.1 mm) coupled to a Waters Acquity C18 analytical column (100 × 2.1 mm × 1.7 µm particles) performed the chromatography process. A mobile phase of 1.0% (v/v) Acetic acid in water (Solvent A) and Acetonitrile (Solvent B) was operated by the UPLC.

The gradient program for elution began with 50% B for 0.5 minutes then 95% B for 4.0 minutes and a 1.0 minute hold. The composition was returned to 50% B at 5.5 minutes followed by the column equilibration for an additional 3.0 minutes. With a flow rate of 0.35 mL and a column held at 35°C, the injection volume was 2µL and the typical Et-IPA retention time was 3.0 minutes. A Waters Xevo TQ triple quadrupole mass spectrometer (Waters Corporation) was joined to the UPLC. Single ion monitoring (SIM) was used simultaneously with multiple reaction monitoring (MRM) in negative electrospray ionisation mode to perform the analyses. The Limit of Quantitation (LOQ) for the method was established in (Pye et al., 2023) as 0.05 ng/mL

### Statistical analysis

Serum Et-IPA concentration values were log transformed prior to analysis. Linear regression analyses were performed using GraphPad Prism v.8.0.

## Results

All individuals received the entire Et-IPA dose in their respective administration methods. We verified the LC-MS/MS analytical method for serum samples from brushtail possums, forester kangaroos and eastern quolls with reference to the original method developed for devil serum samples. No serum extraction samples comprising extraction solution (Methanol:Acetone) alone, along with serum extractions from single (possum) or pooled samples collected from untreated individuals of the same species, confirmed there were no sample matrix interferences (data not shown).

A linear correlation co-efficient of ≥ 0.9995 for each sample was calculated. The external calibration curves showed some species-specific effects of serum as a sample matrix on calibration slopes especially at higher Et-IPA concentration levels (Figure 2). However, substantial sample matrix effects were only observed at serum concentrations well above levels observed in this study. The method performance for the current serum samples was assessed as comparable to that described for the devil study (Pye et al., 2023). Figure 3 shows clear peaks in UPLC-MS/MS chromatograms for all species at retention time 3.0 minutes.

**Figure 3.**
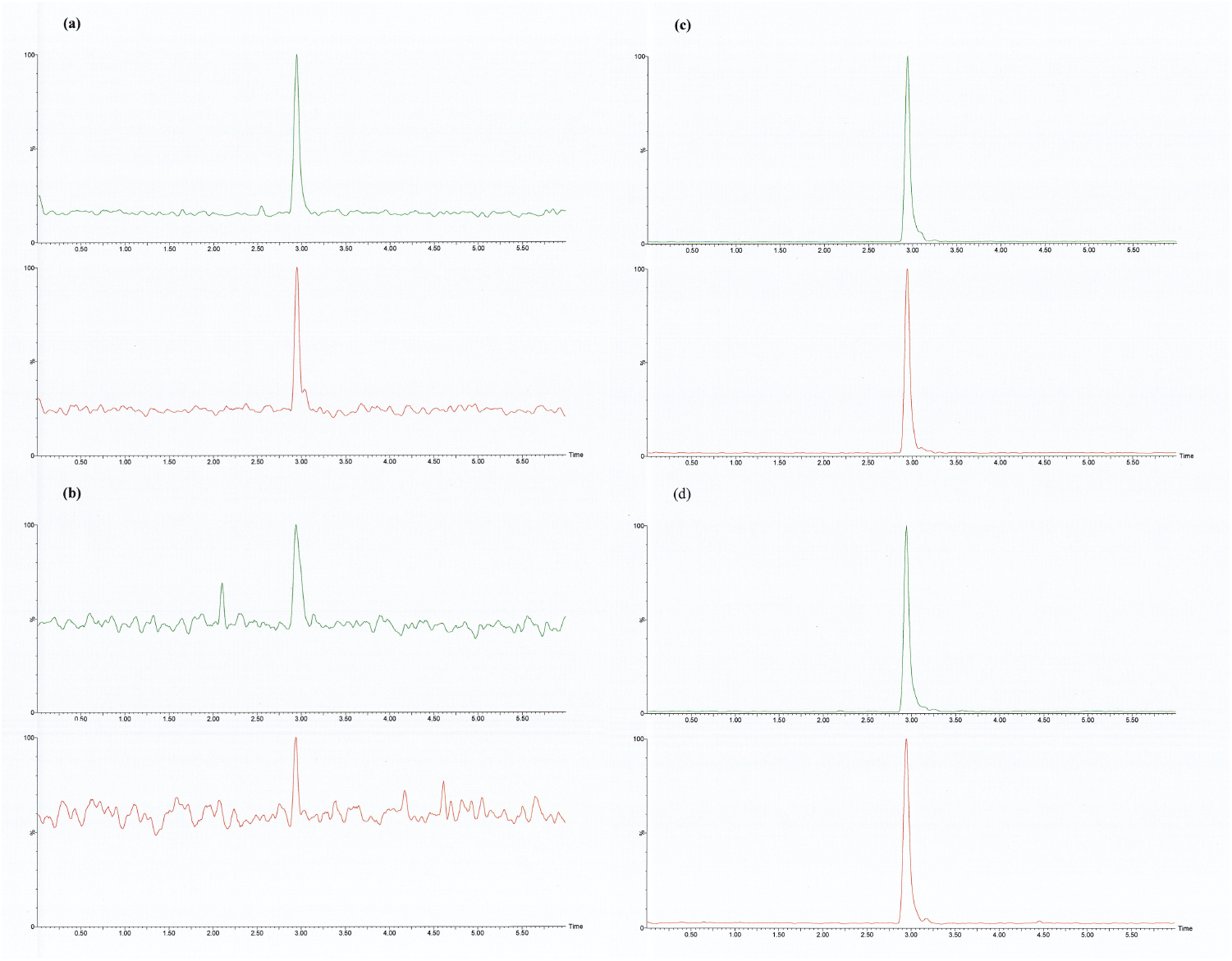
Chromatograms of the three modes of Multiple Reaction Monitoring (MRM) analysis of for detecting Et-IPA in serum samples on day 2 for A: brushtail possum (BTP1), B: forester kangaroo (FK2), C: eastern quoll (EQ1), and D: spotted-tailed quoll (STQ1). Within each figure section, the lower chromatogram represents the MRM transition 1 precursor [M-H]^-^ (m/z) 570.7 to product ion (m/z) 126.8, and the upper chromatogram represents MRM transition 2 precursor [M-H]^-^ (m/z) 570.7 to product ion (m/z) 442.8. For all chromatograms, time in minutes is shown on the x-axis and signal intensity on the y-axis.

Quantifiable serum Et-IPA levels were detected at day two post-consumption in the serum of all individuals in this study for which there was a serum sample (Table 1, Figure 4). One spotted-tailed quoll (STQ2) was unable to be caught on day two for the initial blood collection, and this time point is recorded as not available (NA). On day 14, serum Et-IPA levels were undetectable in all individuals, i.e. below the Limit of Quantitation (LOQ) of 0.05 ng/mL established in (Pye et al., 2024) (Table 1, Figure 4). The proposed blood sample collection on days 28 and 56 were therefore not performed. On day 2, the highest serum Et-IPA concentrations were observed for the species with the highest Et-IPA dose/kg of body weight. Similarly, the lowest serum Et-IPA concentrations were observed for the species with the lowest Et-IPA dose/kg of body weight (Figure 5). Due to small sample sizes and variability in the Et-IPA delivery vehicle, no statistical tests were performed to assess for a potential correlation between dose and body weight.

**Figure 4.**
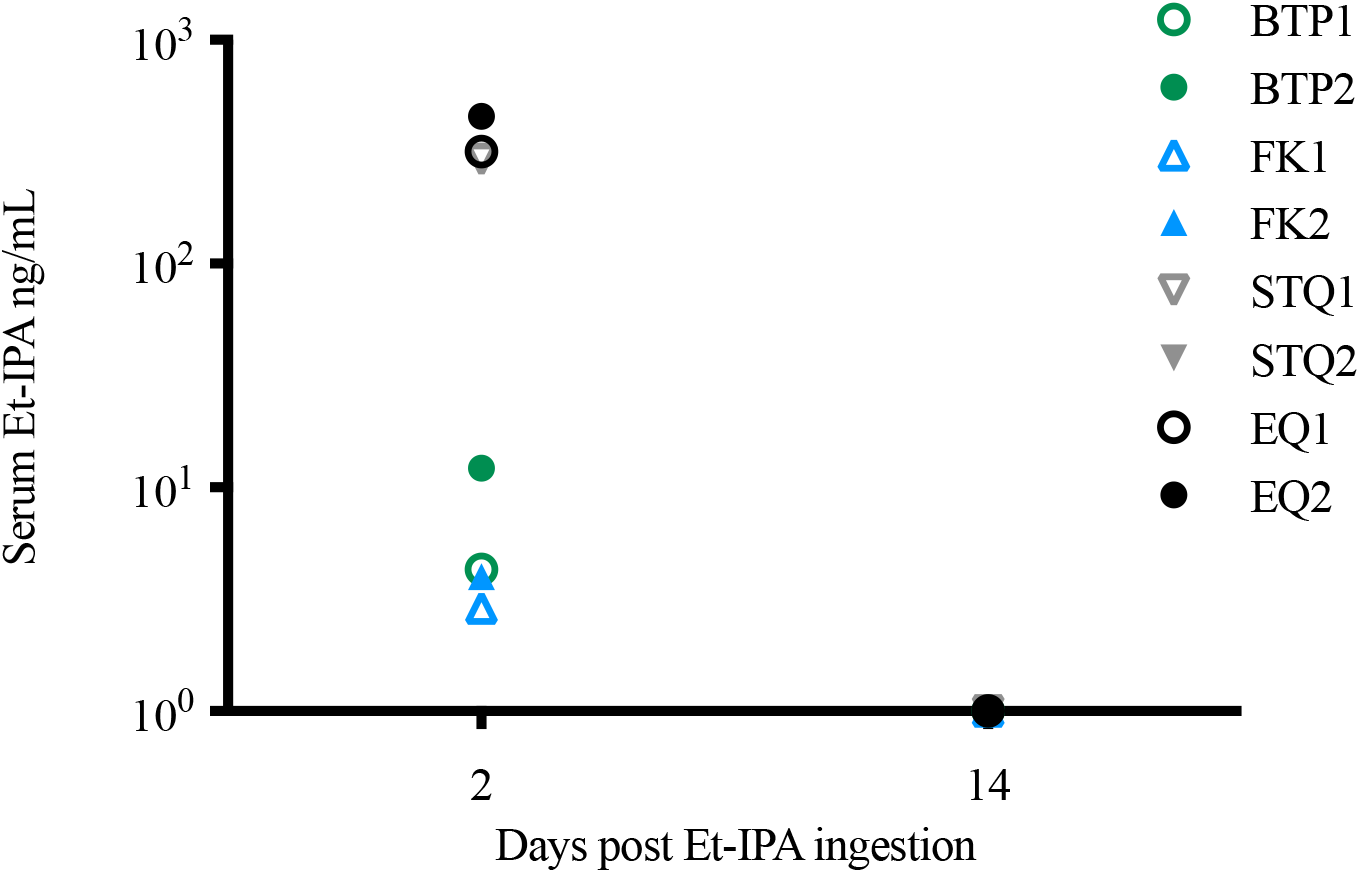
Serum concentration of Et-IPA at day 2 and day 14 post-ingestion of Et-IPA. Serum concentrations were measured in ng/mL and shown on log10 scale. The colour and shape of the symbols indicate the species of each individual. Abbreviations: Brushtail possum 1 (BPT1); brushtail possum 2 (BTP2); forester kangaroo 1 (FK1); forester kangaroo 2 (FK2); spotted-tailed quoll 1 (STQ1); spotted-tailed quoll 2 (STQ2); eastern quoll (EQ1); eastern quoll (EQ2). Result for STQ2 on day 2 not available.

**Figure 5.**
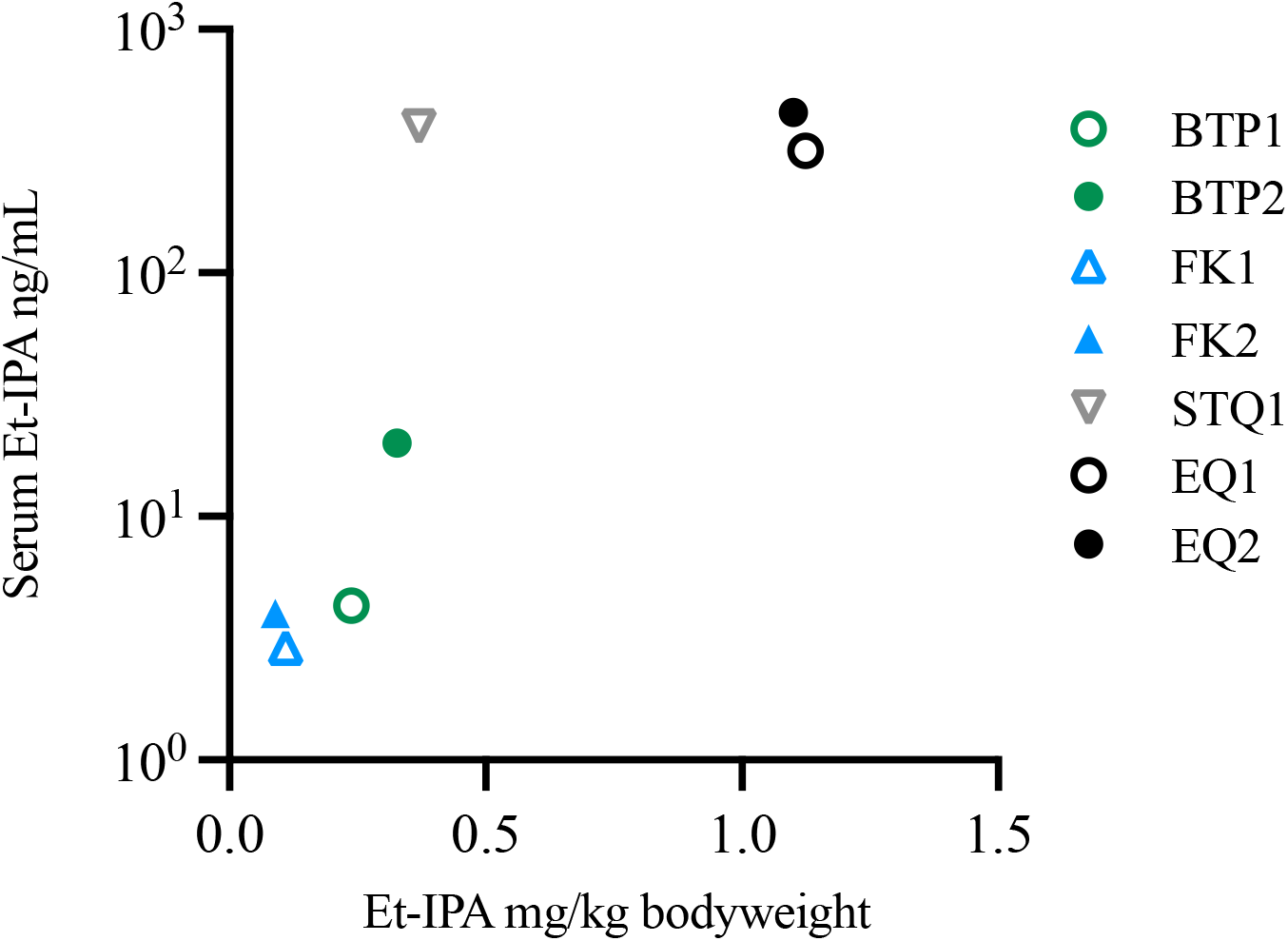
Et-IPA dose and serum Et-IPA concentration for each individual at day 2 post-ingestion. Serum concentration was measured in ng/mL and shown on log_10_ scale. See Figure 4 legend for abbreviations. Result for ST2Q on day 2 not available.

## Discussion

Our results help clarify the contrasting results of previous studies, which suggested that Et-IPA was not a suitable marker for possums (Eason et al., 1994) or wallabies (Fisher and Marks, 1997) but is an effective marker for Tasmanian devils (Pye et al., 2023). The prior studies in possums and wallabies used the less sensitive indirect serum iodine detection method but higher mg/kg doses of Et-IPA and found serum Et-IPA levels decayed to baseline levels in both species by day 10. Our prior study (Pye et al., 2023) found a 1 mg dose of Et-IPA was detectable in the carnivorous marsupial Tasmanian devil for at least 56 days post-ingestion, and for more than 180 days when higher doses were used. Thus, we hypothesized that the more sensitive Et-IPA detection method would be able to detect serum Et-IPA in possums and macropods. In contrast to our hypothesis but in agreement with results of prior studies, the highly sensitive LC-MS/MS method used here did not detect serum Et-IPA in possums or kangaroos by day 14.

We also hypothesised that Et-IPA would be an effective marker for the two quoll species due to their close evolutionary history with devils. The quolls had initial serum Et-IPA levels ranging from 317.7 to 455.3 ng/ml (Table 1). These were notably higher than the possums and kangaroos (2.9 to 20 ng/ml) and approaching the range of serum Et-IPA (560.7 to 1277 ng/ml) detected in devils that were given 1mg Et-IPA in (Pye et al., 2023). In the devil study, individuals were sampled initially on day one while the animals in this study were first sampled on day two due to the feeding behaviours of the eastern quolls as described above in methods. With their apparent rapid elimination of Et-IPA, it is possible the quolls’ results would have been higher on day one. Visual examination of Figure 5 suggests a potential trend for increasing dose per body weight leading to higher serum Et-IPA levels. However, devils are considerably larger than eastern quolls and had much higher serum Et-IPA retention rates. Furthermore, Pye et al., 2023 found the Et-IPA dose was not predictive of serum Et-IPA in devils.

Our studies and those reviewed in Ballesteros et al., 2013 found species administered a wide range of Et-IPA doses vary greatly in the serum marker persistence and decay. Factors affecting the overall fate and clearance of xenobiotics from the body include the binding affinity of albumin and other plasma proteins; metabolism; renal and hepatic excretion mechanisms; and the gut microbiome (Collins and Patterson, 2020, Maxwell, 2024). In the 1950’s, IPA was used as a radiocontrast agent in humans at doses of 3 to 4.5 grams per person, until its exceptionally long half-life of approximately 30 months was realised (Hoffmann, 1954, Ballesteros et al., 2013). This was due in part, if not primarily to the high binding affinity of IPA to human serum albumin (Ryan et al., 2011). Interspecies differences in serum albumin binding affinities have been identified for various compounds (Starnes et al., 2024, Kosa et al., 1997) and likely explain some of the variation in Et-IPA persistence between species. The animals in our study were administered Et-IPA in different foods due to their differing dietary preferences. Other studies have also optimised palatability (e.g. use of peanuts, eggs or foliage as the administration vehicle) to maximise uptake of Et-IPA (Carter et al., 2018, Spurr, 2002, Sweetapple and Nugent, 1999). It is not clear whether different foods affect Et-IPA absorption, however higher serum iodine was found in pigs that had been gavaged when compared to pigs that had consumed the same Et-IPA dose in baits, suggesting that the presence of food might affect absorption (Cowled et al., 2008). The suitability of oily carriers for Et-IPA has been confirmed (Jacoblinnert et al., 2021), and the use of corn oil was consistent across all individuals in our study.

Et-IPA remains a useful biomarker for determining successful bait distribution strategies for delivery of oral vaccines to Tasmanian devils, despite not marking off-target species. While Et-IPA is considered to have a wide margin of safety and low risk of environmental contamination following its excretion in urine or faeces, as reviewed in Ballesteros et al., 2013, minimizing the amount of Et-IPA into the environment is desirable. Bennetts wallabies and Tasmanian pademelons are commercially harvested for human consumption in Tasmania. The subject of secondary uptake of Et-IPA by humans that consumed marked animals was considered by Sage et al., 2013. Their study, in which wild boars (*Sus scrofa*) were administered between 0.9 and 8.4 mg/kg Et-IPA found Et-IPA was detectable in liver and muscle samples from animals tested 217 days after exposure. In our study, all individuals lacked detectable serum Et-IPA two weeks after consumption of the 1 mg dose proposed for optimising devil bait vaccine distribution strategies. This suggests that Et-IPA poses little risk of bioaccumulation in Tasmanian wildlife or humans that consume prey species.

Secondary uptake of Et-IPA via consumption of off-target prey species (macropods and possums) could affect interpretation of results from bait vaccine uptake field trials for Tasmanian devils given that devils opportunistically hunt live macropods and possums and scavenge on their carcasses. However, the rapid elimination of Et-IPA from the four species tested here, along with the low initial serum levels in the prey species suggests that secondary uptake is unlikely to result in bioaccumulation of Et-IPA in devils. More broadly, our study shows that clearance of xenobiotics can vary between species within the same taxonomic family. This reinforces the usefulness of species-specific research for understanding the marking capabilities of Et-IPA in individual species.

