## Supplemental information for "Ethyl-iophenoxic acid as a serum biomarker for marsupial species in oral bait trials"

**Nofs et al 2026**

**Supplementary information**

### Glider porridge

Note from S.A (author) --the amount of glider porridge was in a lid from a cap, approx. 0.1 ml glider porridge.

Brush-Tailed possum given IPA in glider porridge. Ate 100% immediately.

Porridge recipe:
6 x tablespoons of baby cereal (we normally use 'Nestle Cerelac baby rice cereal stage 1)
1.5 x teaspoons of Glucodin powder
1.5 x teaspoons of Sustagen powder (Vanilla)
3 x tablespoons of honey (we normally use 'Gardener mixed blossom honey')
Water – added slowly and mixed with dry ingredients until a 'porridge' like consistency is achieved)
This animal was an outreach animal and trained to take treats on a small dish/lid on a stick.

### Eastern quoll meatball

1 x 50g "quoll ball" each

Eastern Quoll 0.1 ml in quoll meatball. Crated separately with meatball. Left at 4 pm overnight. In morning meatballs w IPA were gone. 100% IPA admin success.
Animals will be separated in crates overnight with their meatball w IPA.

Quoll ball recipe:
2.5kg of minced chicken frames
2x raw eggs (including crushed shell)
250g x rolled oats OR cooked brown rice
50g x InsectaPro powder
250g x wheat bran – used to help take the 'stickiness' out of the mince, more or less may be required in any given batch to achieve the correct consistency)
